# SMC1A facilitates gastric cancer cells proliferation, migration and invasion via promoting SNAI2 activated EMT

**DOI:** 10.1101/2021.09.23.461612

**Authors:** Yaling Liu, Xianrui Fang, Qianqian Wang, Da Xiao, Zhenyu Peng, Ting Zhou, Feng Ren, Jingyu Zhou

## Abstract

**Background:** structural maintenance of chromosomes protein 1A (SMC1A) is a crucial subunit of the cohesion protein complex and plays a vital role in cell cycle regulation, genomic stability maintenance, chromosome dynamics. Recent studies demonstrated that SMC1A participate in tumorigenesis. This reseach aims to explore the role and the underlying mechanisms of SMC1A in gastric cancer (GC).

**Materials and methods:** RT-qPCR and western blot were used to examine the expression levels of SMC1A in GC tissues and cell lines. The role of SMC1A on GC cell proliferation, migration, invasion and epithelial-mesenchymal transition (EMT)were analyzed. Furthermore,the mechanism of SMC1A action was investigated.

**Results:** SMC1A was highly expressed in GC tissues and cell lines. The high expression of SMC1A indicated the poor overall survival of GC patients from Kaplan-Meier Plotter. Enhancing the expression of SMC1A in AGS remarkably promoted cell proliferation, migration and invasion. While knockdown of SMC1A in HCG27 inhibited cell proliferation, migration and invasion of HGC27 cells. Moreover, it’s observed that SMC1A promoted EMT and malignant cell behaviors via regulating SNAI2

**Conclusion:** our study revealed SMC1A facilitates gastric cancer cell proliferation, migration and invasion via promoting SNAI2 activated EMT, which indicated SMC1A may be a potential target for gastric cancer therapy.

## Introduction

Gastric cancer (GC) as one of the most prevalent digestive tract malignancies, seriously damages human health(Liu et al., 2020). Although the incidence has declined in recent years, the mortality rate remains high. The interaction of internal genetic heterogeneity and various external risk factors leads to the occurrence and development of gastric cancer(Karimi et al., 2014). Due to the low detection rate of early screening, and prone to invasion and metastasis of gastric cancer, numerous patients had developed to the middle or late stage when they were diagnosed, and the five-year survival was extremely low(Giordano, 2014). The etiology of gastric cancer is a complicated procedure of multi-gene participation(Liu et al., 2020). Therefore, it’s urgent need to explore an in-depth insight into the underlying mechanisms of GC initiation and development.

Structural maintenance of chromosomes protein 1A (SMC1A), a crucial subunit of the cohesion protein complex, which is complex essential for sister chromatid cohesion in chromosome dynamics(Xiu et al., 2020). SMC1A plays a vital role in cell cycle regulation(Luo et al., 2013), genomic stability maintenance(Diaz-Martinez et al., 2010), chromosome dynamics(Luo et al., 2013, Yan and McKee, 2013, Huber et al., 2016). Recently, numerous studies have reported that SMC1A involves in tumorigenesis(Yi et al., 2017). Such as In prostatic carcinoma, SMC1A was highly expression, and knockdown of SMC1A could not only limited cell proliferation, growth, migration, and cancer stem-like cell properties, but also enhanced efficacy of radiation therapy (Pan et al., 2016, Yadav et al., 2019). Zhang et al (Zhang et al., 2018) reported that phosphorylated SMC1A facilitated hepatocellur carcinoma cell proliferation and migration, meanwhile, the highly expressed of phosphorylated SMC1A was dramatically correlated with worse prognostic outcomes. In colorectal cancers, SMC1A was present as extra-copies, mutations and overexpression, and it contributes to cancer development and metastasis (Sarogni et al., 2019, Zhou et al., 2017). Furthermore, the overexpression of SMC1A indicated as an indepents poor prognostic predictor in late stage of colorectal cancers (Wang et al., 2015). Nevertheless, it was reported that low expression of SMC1A predicts poor survival in acute myeloid leukemia patient(Homme et al., 2010). In gastric cancer, little is known regarding the possible function of SMC1A.

In this research, we analyzed the expession of SMC1A in GC tissues and cell lines, and assessed the correlation between the level of SMC1A and the prognostic survival of GC patients. Furthermore, we evaluated the role and potential mechanism role of SMC1A on gastric cancer cell biology behaviors. Our study sheded light that SMC1A promotes gastric cancer cells proliferation, migration and invasion via promoting SNAI2 activated EMT, this finding may provides a possible therapeutic target for GC.

## Results

### SMC1A was overexpressed in human gastric cancer

To investigate the characteristics in gastric cancer, we analysed SMC1A in the cancer genome atlas (TCGA) stomach adenocarcinoma database. As shown in Figure 1A, the level of SMC1A was evidently increased in the tumor tissues compared with paracancerous tissues (Nomal). In the fresh 20 gastric cancer tissues and the corresponding adjacent tissues, SMC1A expression in cancer tissues was higher than that in corresponding adjacent tissues (Figure 1B). Moreover, the high expression of SMC1A indicated the poor overall survival of GC patients from Kaplan-Meier Plotter <u>(htt</u>p<u>://kmplot.com/analysis/)</u> (Figure 1C). We also examined the expression of SMC1A mRNA and protein in t human gastric cancer cell lines (AGS, HGC27 and NCI-N87) and the human gastric epithelial cell line GES-1. As shown in Figure 1D and 1E, SMC1A was highly expression in GC cell compared wih GES-1.

**Fig 1.**
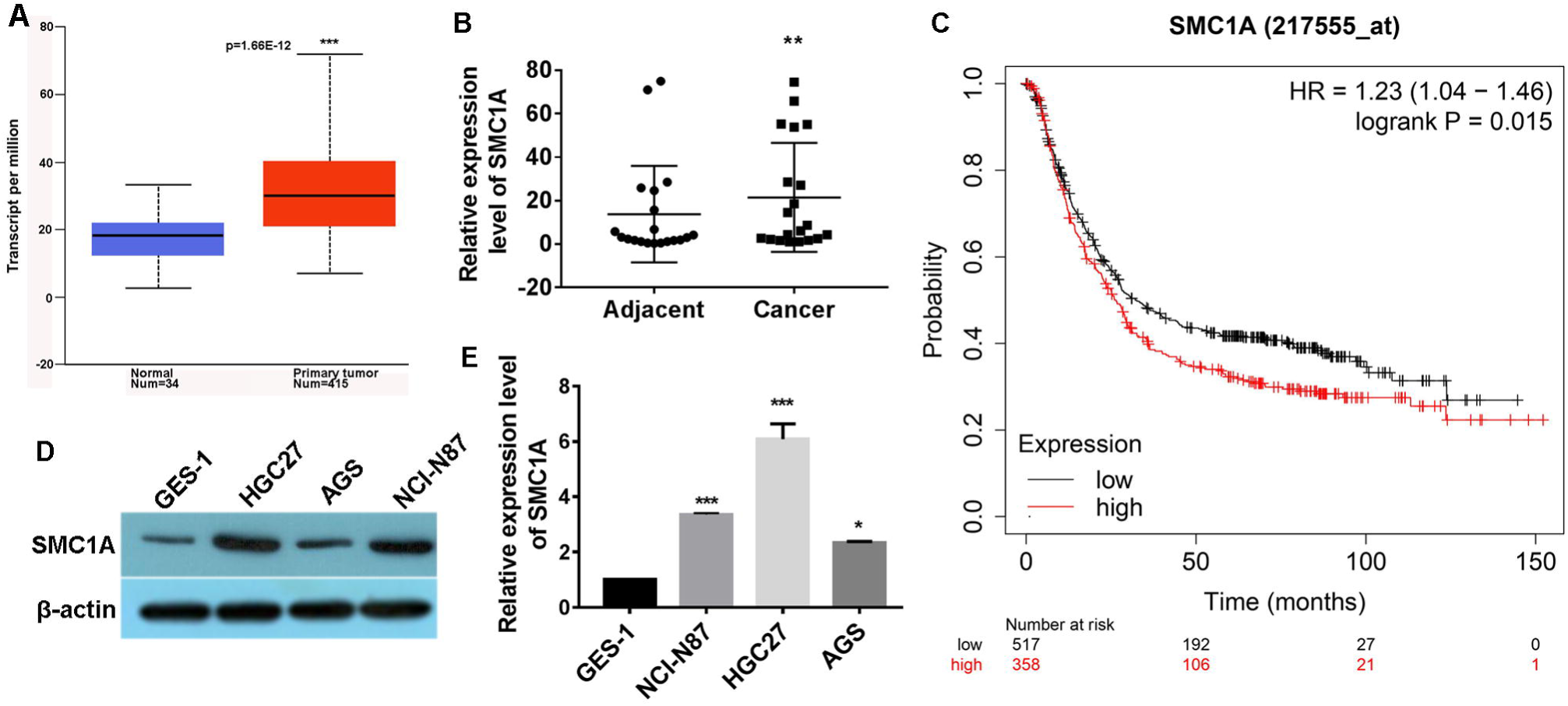
SMC1A was overexpressed in human gastric cancer. (A) The expression of SMC1A was analyzed in cancer tissues from TCGA data. (B) RT-qPCR analysis for SMC1A mRNA in 20 samples of GC tissues and the corresponding adjacent tissues. (C) The overall survival of GC patients was evaluated using Kaplan-Meier Plotter. The expression of SMC1A was examined in human gastric cancer cell lines and the human gastric epithelial cell line GES-1using RT-qPCR method (D) and western blot method (E). **P<0.01, **P<0.001, ***P<0.001.

### SMC1A promoted gastric cancer cell proliferation, invasion and migration

As HGC27 cells had considerably high endogenous SMC1A expression, while AGS cells had low relatively expression, so AGS cells were selected for gain-of-function model by transfecting with SMC1A overexpression plasmid and HGC27 cells was served as loss-of-function model by transfecting SMC1A siRNA. The result of SMC1A knockdown and overexpression was verifitied by western blot method (Figure 2A). CCK-8 assay displayed that the proliferation of HGC27 cells significantly descended after SMC1A depletion, while SMC1A over-expression promoted proliferation of AGS cells (Figure 2B). Colony formation assay confirmed that SMC1A positively regulated colony formation ability in HGC27 and AGS cells (Figure 2C). Transwell invasion showed that SMC1A depletion decreased the invasion in HGC27 cells, while SMC1A over-expression increased AGS cells invasion (Figure 2D). Wound healing assay demonstrated that SMC1A over-expression promoted AGS cells migration. while SMC1A knockdown inhibited HGC27 cell migration. (Figure 2E). It’s suggested that SMC1A may promoted GC cell proliferation, migration and invasion.

**Fig 2.**
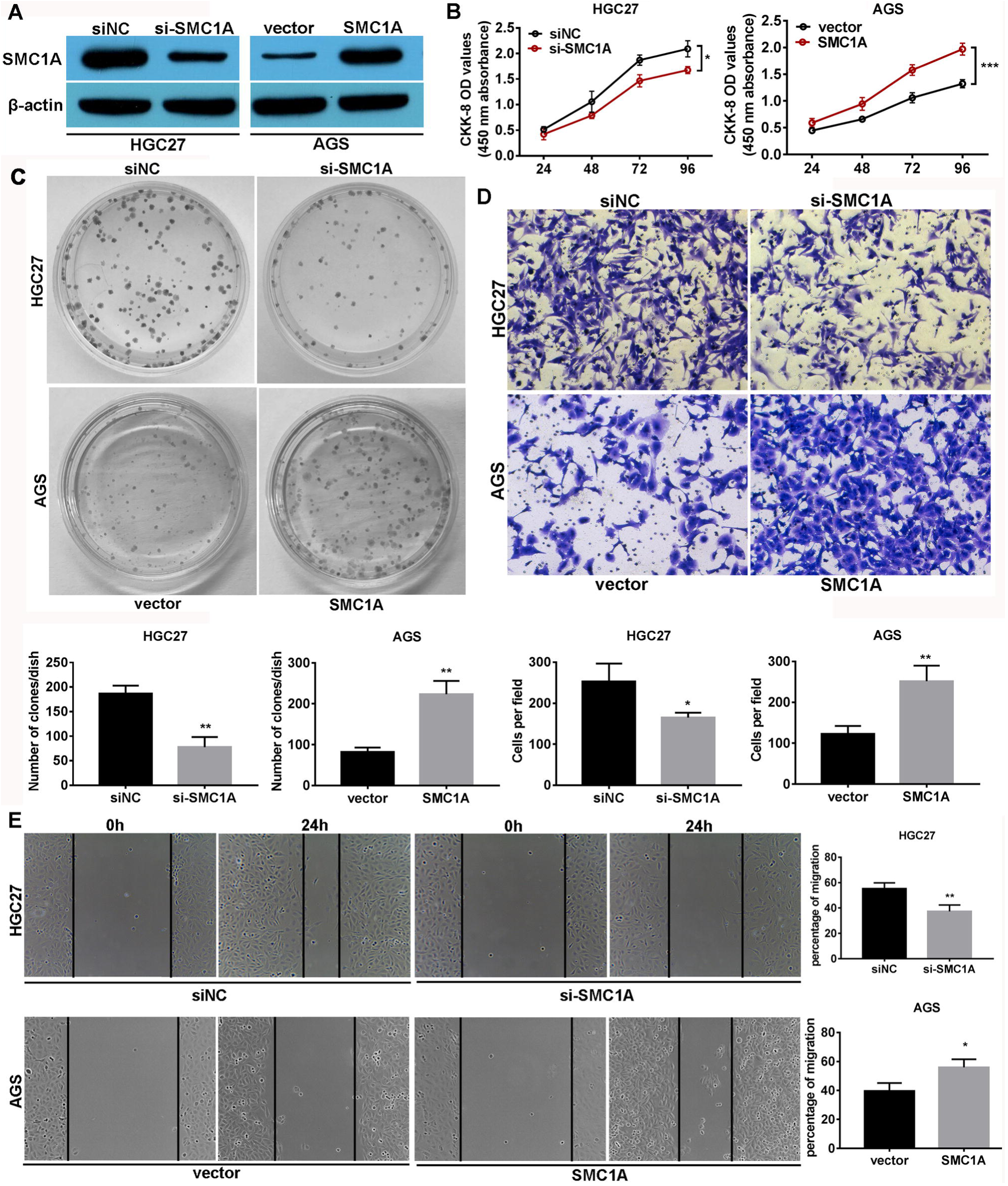
SMC1A gastric cancer cell proliferation, invasion and migration. (A) The expression of SMC1A was examined in SMC1A silenced and overexpressed cells by western blot method. CCK-8 assay (B) and Colony formation assay (C) were used to determine the effect of SMC1A knockdown and overexpression on cell proliferation. Matrigel invasion assay (D) and Wound healing assay (F) analyzed the effect of SMC1A knockdown and overexpression on cell invasion and migration respectively. *P<0.05, **P<0.01, ***P<0.001.

### SMC1A promoted EMT via upregulating SNAI2

Epithelial-mesenchymal transition (EMT) is a process which epithelial cells tansform inton a mesenchymal phenotype, concomitantly reduce the expression of E-cadherin regulated by one or several factors such as; SNAIL, SLUG, ZEBs and TWIST (Hardin et al., 2018). Notably, through Chip assay, Noutsou et al (Noutsou et al., 2017) have demonstrated that SMC1A promotes progenitor function by maintaining open chromatin to upregulate the expression of self-renewal gene SNAI2. Thus, we speculated that SMCIA may be involved in EMT process via regulation SNAI2. As showing in Figure 3A, SMC1A knockdown obviously decreased SNAI2 expression, while SMC1A overexpression elevated SNAI2 levle. The detectation of EMT related marker showed that the expression of mesenchymal markers N-cadherin and Vimentin was reduced and epithelial marker E-cadherin was increased in SMC1A depletion cells. Converse results were observed in the SMC1A overexpression cells (Figure 3B). Besides, the similar results were seen by immunofluorescence assay (Figure 3C and 3D). Furthermore, restoring SNAIL2 expression in SMC1A silenced cells could significant mitigated the expression change of EMT related markers (Figure 3E and 3F). These results implied that SMC1A facilitated EMT via upregulating SNAI2

**Fig 3.**
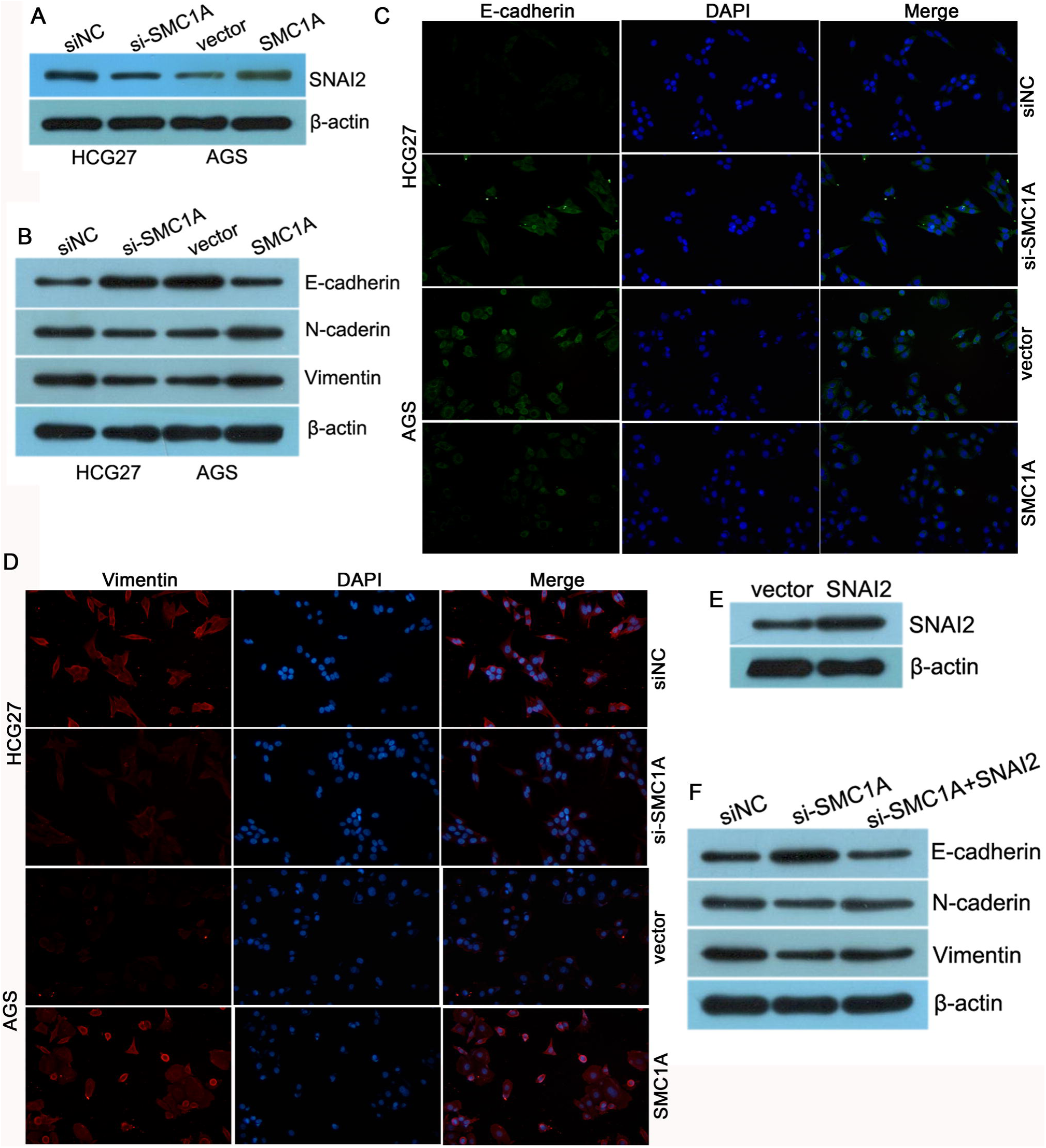
SMC1A promoted GC epithelial-mesenchymal transition (EMT) via upregulating SNAI2. (A) SNAI2 level was examined in SMC1A silenced and overexpressed cells using western blot method. (B) Proteins level of EMT markers E-cadherin, N-cadherin and Vimentin were detected after SMC1A depletion and overexpression. Immunofluorescence assay detected the expression of E-cadherin (C) and Vimentin (D) in SMC1A silenced and overexpressed cells. (E) The expression of SNAI2 was examined in SNAI2 overexpressed cells by western blot method. (F) Proteins level of EMT markers E-cadherin, N-cadherin and Vimentin were detected in response to the treatment of SMC1A siRNA and SMC1A siRNA+SNAI2.

### SMC1A promoted GC cell proliferation, invasion and migration via SNAI2

Numerous literature have been reported that EMT may promotes cell growth, migration and metastatic in GC (Bure et al., 2019, Landeros et al., 2020, Yue et al., 2019). Therefore, rescue experiments were performed to evaluate whether SMC1A promoted GC biology behaviors via EMT regulator SNAI2. As determined by CCK-8, clone formation, Transwell invasion and wound-healing assays, restoring SNAI2 expression could evidently attenuated the suppressive effect on GC cell proliferation, clone formation, migration and invasion causing by SMC1A silencing (Figure 4A-D). It’s indicated that SMC1A promoted GC cell proliferation, migration and invasion via SNAI2.

**Fig 4.**
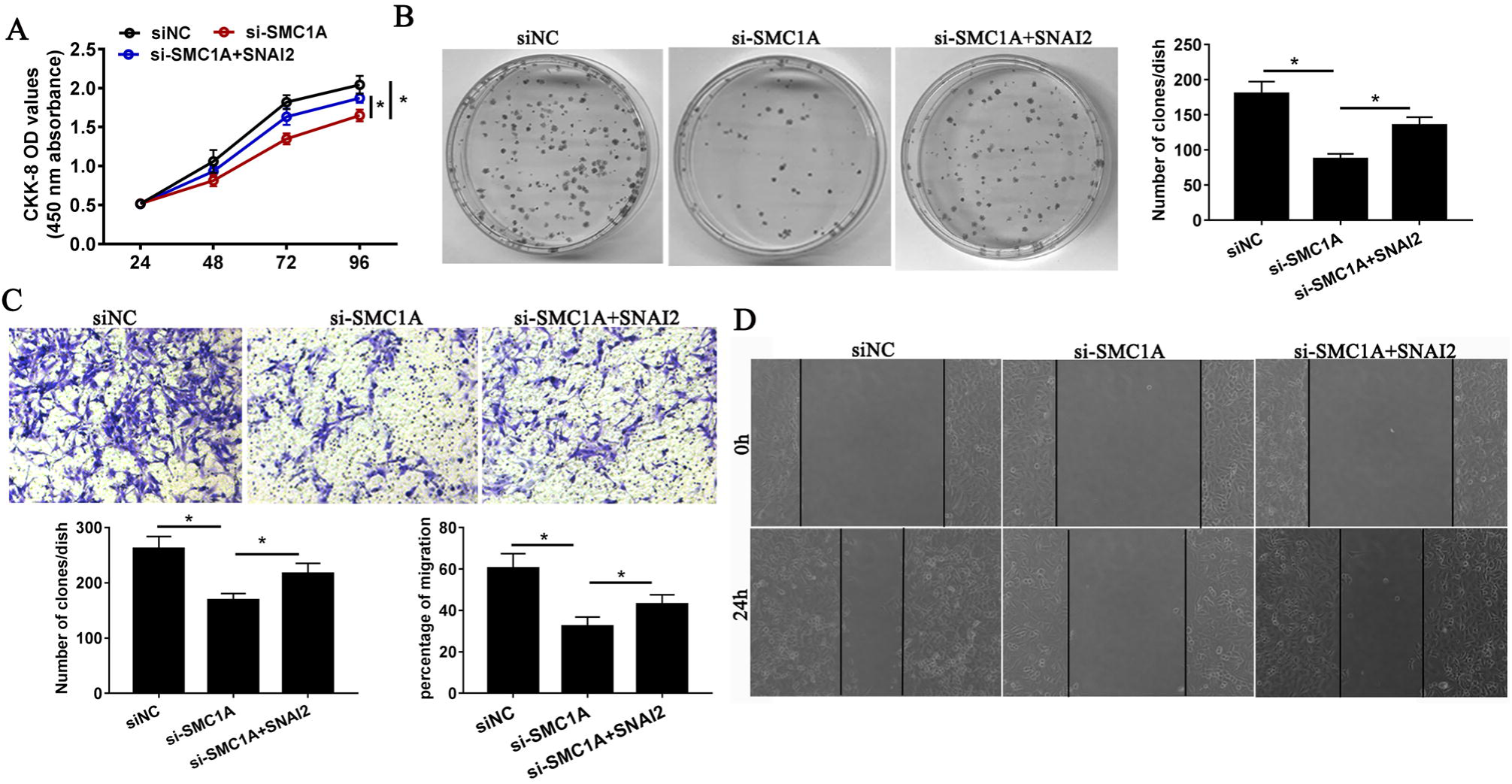
SMC1A promoted GC cell proliferation, invasion and migration via SNAI2. CCK-8 assay (A) and Colony formation assay (B) were performed to analysis cell proliferation in response to the treatment of SMC1A siRNA and SMC1A siRNA+SNAI2. Matrigel invasion assay (C) and Wound healing assay (D) used to investigate cell invasion and migration in response to the treatment of SMC1A siRNA and SMC1A siRNA+SNAI2. *P<0.05.

## Disscussion

Gastric cancer influences nearly 2 million peoples every year, 75–85% of whom die within 5 years after diagnosis, which making it the third most fatal cancer worldwide(Oliveira et al., 2015). Currently, lack of insufficient diagnoses at early stage and the characterinstics of tumor prone to invasion and metastasis were a thorny problem in GC treatment. Although the treatments such as chemotherapy, radiotherapy, palliative resection and gastrectomy have made significant improvement, the prognosis were not encouraging (Li et al., 2020). Thus, it’s necessary to explore the underlying molecular mechanism mechanisms of GC initiation and development, which will contributes to provide treatment strategies and monitor the prognosis of GC. In this study, SMC1A was upregulation in GC tumor tissues and cells, and the high expression of SMC1A associated with the poor overall survival. Cell function and mechanism assays revealed that SMC1A facilitates gastric cancer cells proliferation, migration and invasion via promoting SNAI2 activated EMT.

As an important structural maintenance factor of chromosome proteins, SMC1A participates in many biological processes(Zhang et al., 2018). Previous research had revealed that SMC1A plays an important role in tumorigenesis and progressions (Yi et al., 2017). In most of tumors, SMC1A was reported as a proto-oncogene that could promoted cancer development and metastasis (Pan et al., 2016, Zhang et al., 2018, Zhou et al., 2017). But in acute myeloid leukemia, SMC1A was downregulation and its low expression predicts poor survival, SMC1A overexpression could inhibited cell proliferation and promoted apoptosis (Homme et al., 2010, Zhao et al., 2019). Consistent with most literature, our study found that SMC1A could promoted gastric cancer cell proliferation, migration and invasion in vitro. Howerver, the detailed molecular mechanism of SMC1A exerting in cancers had no reported so far.

Epithelial-mesenchymal transition (EMT) was first reported in embryonic development and played a vital role in the formation of germ layer and organs (Zhou et al., 2019). During EMT process, epithelial cells miss the polarity and intrecellular adhesion, and acquire various of mesenchymal phenotype, as motility and invasive (Ye and Weinberg, 2015). This process was induced by several EMT associated factors, such as the epithelial marker E-cadherin, the mesenchymal markers N-cadherin and vimentin, and the group of transcriptional repressors Zeb, Snail, Slug and Twist (Cao et al., 2020b). Accumulated evidences have revealed that EMT participates in tumor invasion and metastasis (Cao et al., 2020a, Maruta and Greer, 1988, Zhou et al., 2019). The reducing of E-caherin expression is one of significant EMT hallmark, as it’s the crucial epithelial marker of EMT which distrubuted in epithelial cells junctions and participates in the composition of intercellular adhesion complex (Zhou et al., 2019). Once E-cadherin is lose, causing the loose of tight links among cells. Simutaneously, cells adhesion is reduced, which making cells facilitates to detach from the primary tumor site, and enhances cells exercise and invasion capacity (MacDonald et al., 2009, Zhou et al., 2019). Previous study has been demonstrated that SMC1A mediated raioresistance in prostate cancer by regulating EMT (Yadav et al., 2019). In our study, we also observed SMC1A could induced EMT in gastric cancer. But the mechanism of SMC1A modulation EMT has not been reported. Snail family transcriptional repressor 2 (SNIL2) also named slug, as a crucial EMT inucible factor, which acted as the transcriptional suppressor of E-cadherin via binding to the conserved E-box motif in the promoter region (Chen et al., 2019). It has been reported that SNAI2 could promotes GC cell EMT, invasion and metastasis (Dong et al., 2021, Sakimura et al., 2015, Zhou et al., 2019). In this resech, we found that SMC1A could promoted EMT and malignant cell behaviors via regulating SNAI2, which provides a good sight in understaning the oncogenic role of SMC1A in GC.

In summary, our study revealed SMC1A facilitates gastric cancer cell proliferation, migration and invasion via promoting SNAI2 activated EMT, which indicated SMC1A may be a potential target for gastric cancer therapy

## Materials and methods

### Clinical specimens

A total of 20 primary stomach adenocarcinoma cancer tissues (n=20) and their matched adjacent normal tissues (n=20) were collected from the Department of Geriatric Surgery, the Second Xiangya Hospital, Central South University from 2018 to 2019. No patient receive chemotherapy or radiotherapy before surgical resection. This study was approved by the Ethics Committee of the Second Xiangya Hospital of Central South University. All the patients have signed informed consent. The Ethics Committee approved. Collection of tissue samples. The fresh tissues were fast frozen in liqud nitrogen and stored at -80°C until use.

### Cell culture and transfection

Human gastric cancer cells AGS and HGC27 were acquired from Cell Bank of Chinese Academy of Sciences (Shanghai, China). AGS cells were culture in F12K medium, while HGC27 cells were maintained in RPMI 1640 medium. Two mediums were supplemented with 10% fetal bovine serum(FBS) (Gibco, USA). All cells were cultured at 37 °C in a humidified incubator with 5% CO_2_.

The small interfering RNA (siRNA) of homo SMC1A were designed and purchased from General Biosystems (Anhui, China). siRNA and the negative control siRNA (siNC) with 20 nM in 6-well plates were transfected using Liopfectamine 3000 (Invitrogen, Carlsbad, USA) according to the manufacturers’ protocols. Total RNA and protein were extracted after a 48 h transfection. The interference sequences were shown in Table 1.

**Table1.**
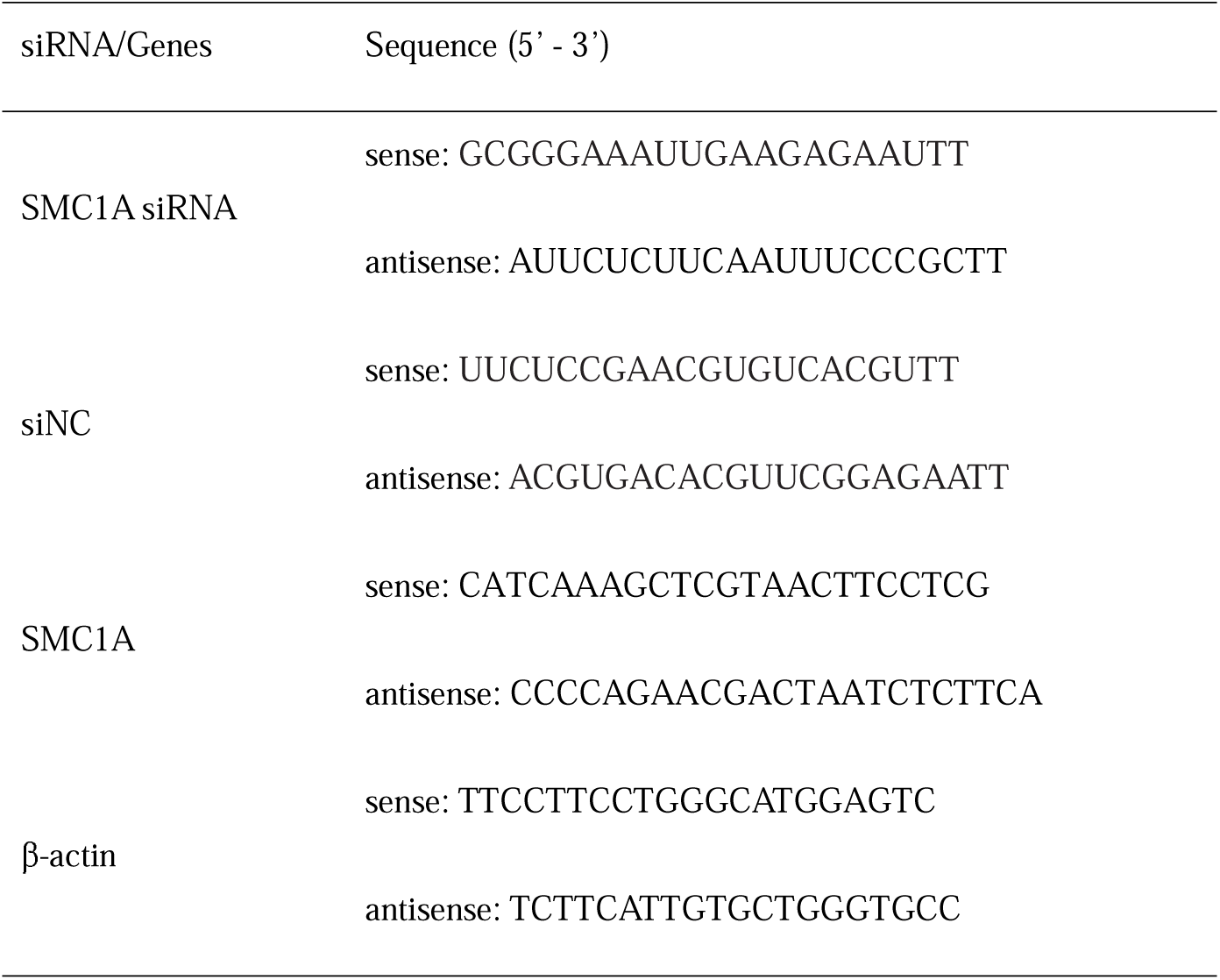
The sequence information of siRNAs and primers.

The SMC1A cDNA ORF clone (HG18194-UT) and SNAI2 cDNA ORF clone (HG11196-M) were purchased from SinoBiological(Beijing, China), and subcloned into a pCDNA3.1 vector. Plasmids were transfected into cells using Lipofectamine 3000 (Thermo Fisher, USA) following the manufacturers’ instructions. Cells were harvested at 48 h after transfection.

### RNA isolation and reverse transcription-quantitative PCR(RT-qPCR)

Total RNA was isolated using TRIzol Reagent (TIANGEN, Beijing, China). Reverse transcription was executed using the Revertaid First Strand cDNA Synthesis Kit (Thermo Fish, Carlsbad, CA, USA) following the manufacturer’s instructions. The expression of mRNAs were quantified by RT-qPCR analysis with SYBR Green PCR Kit(Invitrogen, California, USA). β-actin was used for the normalization control, and calculated based on the 2^−ΔΔCT^ method. The sequences of primers were shown in Table 1.

### Western blot analysis

Total proteins were isolated from GC cell and the concentration was measured with BCA Kit (ThermoFisher, USA). Proteins were separated by 10% SDS-PAGE gel and transferred to 0.45 µm polyvinylidene difluoride (PVDF) membrane (Millipore, USA). After blocking by with 5% non-fat dry milk for 2h at room temperature, The membrances were incubated with primary antibodies against SMC1A (1:2000, Immunoway), SNAI2 (1:1000, CST), E-cadherin (1:2000, Proteintech), N-cadherin(1:1000, Proteintech), Vimentin (1:2000, Proteintech) overnight at 4°C, followed by incubating with HRP-labeled secondary antibody. The bloting was visualized using enhanced chemiluminescence reagent (Thermofisher, USA). β-actin used as control.

### Cell proliferation assay

Cell proliferation ability was evaluated by Cell Counting Kit-8 (CCK-8) (Beyotime, China) assays. AGS or HGC27 cells were seeded separately into 96-well plates with the density of 4 × 10^3^ cells/well. The second day, the medium supernatant was removed. Subsequently, 100 µl basic RPIM 1640 medium or basic F12K medium were added into HGC27 and AGS cells with 10 µl CCK8, respectively. The cells were incubated for another 2 h. Ultimately, the optical density was measured by microplate Reader (Victor3 1420 Multilabel Counter, Perkin Elmer, USA) at 450 nm.

### Colony formation assay

AGS or HGC27 cells were implanted in 6-well plates (1000 cells/well, approximately). Then renewed the medium. After growing for another 14 days, the cells were washed triple with phosphate buffer, fixed in 4% paraformaldehyde (Solarbio, Beijing, China) 30 min, and then stained with 0.5% crystal violet (Solarbio, Beijing, China).

### Cell invasion assay

Cell invasion assays were performed in a transwell chamber coverd with Matrigel (BD Biosicences, Bedford, USA). After 48 h of transfection, 1×10^5^ AGS or HGC27 cells were plated in the upper chambers with serum-free medium. By contraries, medium with 20% FBS were added into the lower chambers. After 48 hours incubation, the invading cells were fixed in 4% paraformaldehyde and stained with 0.5% crystal viole.

### Wound healing assay

AGS and HGC27 cells were cultured in 6-well plates. When cells grew to about 80% confluence, cells were scratched by 10 µl plastic pipette tip. Next, cells were washed with PBS to remove debris. The wound-healing distance was measured at the 0 h and after 24 h to statistical cell migration.

### Immunofluorescence

Cells on sterile slips were fixed in 4% parafomaldehyde for 15 min and permeabilized with 0.1% Triton X-100 for 10 min at room temperature. Next, the slips were incubated with E-cadherin (1:50, #20874-1-AP, Proteintech) or Vimentin (1:100, #60330-1-Ig, Proteintech) antibody overnight at 4 °C. Subsequently, the slips were incubated with FITC labeled goat anti-rabbit IgG (1:1000, #ab6717, Abcam) or Cy3-AffiniPure Goat Anti-Mouse IgG (1:500, #115-165-003, Jackson) for 1 hours. DAPI was used for nuclear staining. Finally, fluorescence was imaged under the fluorescent microscope (IX71, Olympus, Japan).

### Statistical analysis

Data are analyzed by the GraphPad Prism 7.0. The difference among groups was determined by T-test. All the experiments were repeated at least three timens. *P*<0.05 was considered as statistically significant.

## References

Bure, I. V., Nemtsova, M. V. & Zaletaev, D.V. 2019. Roles of E-cadherin and Noncoding RNAs in the Epithelial-mesenchymal Transition and Progression in Gastric Cancer. Int J Mol Sci, 20.

Cao, R., Ma, B., Wang, G., Xiong, Y., Tian, Y. & Yuan, L. 2020a. An Epithelial-Mesenchymal Transition (EMT) Preoperative Nomogram for Prediction of Lymph Node Metastasis in Bladder Cancer (BLCA). Dis Markers, 2020, 8833972.

Cao, R., Yuan, L., Ma, B., Wang, G., Qiu, W. & Tian, Y. 2020b. An EMT-related gene signature for the prognosis of human bladder cancer. J Cell Mol Med, 24, 605–617.

Chen, D. D., Cheng, J. T., Chandoo, A., Sun, X. W., Zhang, L., Lu, M. D., Sun, W. J. & Huang, Y. P. 2019. microRNA-33a prevents epithelial-mesenchymal transition, invasion, and metastasis of gastric cancer cells through the Snail/Slug pathway. Am J Physiol Gastrointest Liver Physiol, 317, G147–G160.

Diaz-Martinez, L. A., Beauchene, N. A., Furniss, K., Esponda, P., Gimenez-Abian, J. F. & Clarke, D.J. 2010. Cohesin is needed for bipolar mitosis in human cells. Cell Cycle, 9, 1764–73.

Dong, D., Na, L., Zhou, K., Wang, Z., Sun, Y., Zheng, Q., Gao, J., Zhao, C. & Wang, W. 2021. FZD5 prevents epithelial-mesenchymal transition in gastric cancer. Cell Commun Signal, 19, 21.

Giordano, T. J. 2014. The cancer genome atlas research network: a sight to behold. Endocr Pathol, 25, 362–5.

Hardin, H., Helein, H., Meyer, K., Robertson, S., Zhang, R., Zhong, W. & Lloyd, R.V. 2018. Thyroid cancer stem-like cell exosomes: regulation of EMT via transfer of lncRNAs. Lab Invest, 98, 1133–1142.

Homme, C., Krug, U., Tidow, N., Schulte, B., Kuhler, G., Serve, H., Burger, H., Berdel, W. E., Dugas, M., Heinecke, A., Buchner, T., Koschmieder, S. & Muller-Tidow, C. 2010. Low SMC1A protein expression predicts poor survival in acute myeloid leukemia. Oncol Rep, 24, 47–56.

Huber, R. G., Kulemzina, I., Ang, K., Chavda, A. P., Suranthran, S., Teh, J. T., Kenanov, D., Liu, G., Rancati, G., Szmyd, R., Kaldis, P., Bond, P. J. & Ivanov, D. 2016. Impairing Cohesin Smc1/3 Head Engagement Compensates for the Lack of Eco1 Function. Structure, 24, 1991–1999.

Karimi, P., Islami, F., Anandasabapathy, S., Freedman, N. D. & Kamangar, F. 2014. Gastric cancer: descriptive epidemiology, risk factors, screening, and prevention. Cancer Epidemiol Biomarkers Prev, 23, 700–13.

Landeros, N., Santoro, P. M., Carrasco-Avino, G. & Corvalan, A.H. 2020. Competing Endogenous RNA Networks in the Epithelial to Mesenchymal Transition in Diffuse-Type of Gastric Cancer. Cancers (Basel), 12.

Li, P., Wang, L., Li, P., Hu, F., Cao, Y., Tang, D., Ye, G., Li, H. & Wang, D. 2020. Silencing lncRNA XIST exhibits antiproliferative and proapoptotic effects on gastric cancer cells by up-regulating microRNA-132 and down-regulating PXN. Aging (Albany NY), 13, 14469–14481.

Liu, T., Yang, S., Sui, J., Xu, S. Y., Cheng, Y. P., Shen, B., Zhang, Y., Zhang, X. M., Yin, L. H., Pu, Y. P. & Liang, G.Y. 2020. Dysregulated N6-methyladenosine methylation writer METTL3 contributes to the proliferation and migration of gastric cancer. J Cell Physiol, 235, 548–562.

Luo, Y., Deng, X., Cheng, F., Li, Y. & Qiu, J. 2013. SMC1-mediated intra-S-phase arrest facilitates bocavirus DNA replication. J Virol, 87, 4017–32.

Macdonald, B. T., Tamai, K. & He, X. 2009. Wnt/beta-catenin signaling: components, mechanisms, and diseases. Dev Cell, 17, 9–26.

Maruta, S. & Greer, M.A. 1988. Evidence that thyroxine inhibits either basal or TRH-induced TSH secretion only after conversion to triiodothyronine. Proc Soc Exp Biol Med, 187, 391–7.

Noutsou, M., Li, J., Ling, J., Jones, J., Wang, Y., Chen, Y. & Sen, G.L. 2017. The Cohesin Complex Is Necessary for Epidermal Progenitor Cell Function through Maintenance of Self-Renewal Genes. Cell Rep, 20, 3005–3013.

Oliveira, C., Pinheiro, H., Figueiredo, J., Seruca, R. & Carneiro, F. 2015. Familial gastric cancer: genetic susceptibility, pathology, and implications for management. Lancet Oncol, 16, e60–70.

Pan, X. W., Gan, S. S., Ye, J. Q., Fan, Y. H., Hong, U., Chu, C. M., Gao, Y., Li, L., Liu, X., Chen, L., Huang, Y., Xu, H., Ren, J. Z., Yin, L., Qu, F. J., Huang, H., Cui, X. G. & Xu, D.F. 2016. SMC1A promotes growth and migration of prostate cancer in vitro and in vivo. Int J Oncol, 49, 1963–1972.

Sakimura, S., Sugimachi, K., Kurashige, J., Ueda, M., Hirata, H., Nambara, S., Komatsu, H., Saito, T., Takano, Y., Uchi, R., Sakimura, E., Shinden, Y., Iguchi, T., Eguchi, H., Oba, Y., Hoka, S. & Mimori, K. 2015. The miR-506-Induced Epithelial-Mesenchymal Transition is Involved in Poor Prognosis for Patients with Gastric Cancer. Ann Surg Oncol, 22 Suppl 3, S1436–43.

Sarogni, P., Palumbo, O., Servadio, A., Astigiano, S., D’alessio, B., Gatti, V., Cukrov, D., Baldari, S., Pallotta, M. M., Aretini, P., Dell’Orletta, F., Soddu, S., Carella, M., Toietta, G., Barbieri, O., Fontanini, G. & Musio, A. 2019. Overexpression of the cohesin-core subunit SMC1A contributes to colorectal cancer development. J Exp Clin Cancer Res, 38, 108.

Wang, J., Yu, S., Cui, L., Wang, W., Li, J., Wang, K. & Lao, X. 2015. Role of SMC1A overexpression as a predictor of poor prognosis in late stage colorectal cancer. BMC Cancer, 15, 90.

Xiu, D., Liu, L., Cheng, M., Sun, X. & Ma, X. 2020. Knockdown of lncRNA TUG1 Enhances Radiosensitivity of Prostate Cancer via the TUG1/miR-139-5p/SMC1A Axis. Onco Targets Ther, 13, 2319–2331.

Yadav, S., Kowolik, C. M., Lin, M., Zuro, D., Hui, S. K., Riggs, A. D. & Horne, D.A. 2019. SMC1A is associated with radioresistance in prostate cancer and acts by regulating epithelial-mesenchymal transition and cancer stem-like properties. Mol Carcinog, 58, 113–125.

Yan, R. & Mckee, B.D. 2013. The cohesion protein SOLO associates with SMC1 and is required for synapsis, recombination, homolog bias and cohesion and pairing of centromeres in Drosophila Meiosis. PLoS Genet, 9, e1003637.

Ye, X. & Weinberg, R.A. 2015. Epithelial-Mesenchymal Plasticity: A Central Regulator of Cancer Progression. Trends Cell Biol, 25, 675–686.

Yi, F., Wang, Z., Liu, J., Zhang, Y., Wang, Z., Xu, H., Li, X., Bai, N., Cao, L. & Song, X. 2017. Structural Maintenance of Chromosomes protein 1: Role in Genome Stability and Tumorigenesis. Int J Biol Sci, 13, 1092–1099.

Yue, B., Song, C., Yang, L., Cui, R., Cheng, X., Zhang, Z. & Zhao, G. 2019. METTL3-mediated N6-methyladenosine modification is critical for epithelial-mesenchymal transition and metastasis of gastric cancer. Mol Cancer, 18, 142.

Zhang, Y., Yi, F., Wang, L., Wang, Z., Zhang, N., Wang, Z., Li, Z., Song, X., Wei, S. & Cao, L. 2018. Phosphorylation of SMC1A promotes hepatocellular carcinoma cell proliferation and migration. Int J Biol Sci, 14, 1081–1089.

Zhao, C., Wang, S., Zhao, Y., Du, F., Wang, W., Lv, P. & Qi, L. 2019. Long noncoding RNA NEAT1 modulates cell proliferation and apoptosis by regulating miR-23a-3p/SMC1A in acute myeloid leukemia. J Cell Physiol, 234, 6161–6172.

Zhou, P., Xiao, N., Wang, J., Wang, Z., Zheng, S., Shan, S., Wang, J., Du, J. & Wang, J. 2017. SMC1A recruits tumor-associated-fibroblasts (TAFs) and promotes colorectal cancer metastasis. Cancer Lett, 385, 39–45.

Zhou, Q., Chen, S., Lu, M., Luo, Y., Wang, G., Xiao, Y., Ju, L. & Wang, X. 2019. EFEMP2 suppresses epithelial-mesenchymal transition via Wnt/beta-catenin signaling pathway in human bladder cancer. Int J Biol Sci, 15, 2139–2155.

